# Saliva metabolome alterations after acute stress

**DOI:** 10.1101/2022.02.24.480641

**Authors:** Liat Morgan, Rune Isak Dupont Birkler, Shira Shaham-Niv, Yonghui Dong, Tal Wachsman, Lior Carmi, Boris Yakobson, Lihi Adler-Abramovich, Hagit Cohen, Joseph Zohar, Melissa Bateson, Ehud Gazit

## Abstract

Major stress has systemic effects on the body that can have adverse consequences for physical and mental health. However, the molecular basis of these damaging effects remains incompletely understood. Here we use a longitudinal approach to characterise the acute systemic impact of major psychological stress in a pig model. We perform untargeted metabolomics on non-invasively-obtained saliva samples from pigs before and 24-hours after transfer to the novel physical and social environment of a slaughterhouse. The main molecular changes occurring include decreases in amino acids, B-vitamins, and amino acid-derived metabolites synthesized in B-vitamin-dependent reactions, as well as yet-unidentified metabolite features. Decreased levels of several of the identified metabolites are implicated in the pathology of human psychological disorders and neurodegenerative disease, suggesting a possible neuroprotective function. Our results provide a fingerprint of the acute effect of psychological stress on the metabolome and suggest candidate biomarkers with potential roles in stress-related disorders.

**One Sentence Summary:** Identification of metabolites decreased under acute psychological stress may lead to future discoveries for the prevention and treatment of stress-related disorders.

## INTRODUCTION

Psychological stress has systemic effects on the body that may lead to chronic impairments in physical and mental health, resulting in reductions in longevity and quality of life (Schneiderman, Ironson and Siegel, 2005; Yaribeygi *et al*., 2017). More than 70% of human diseases are considered to be stress-related (Lee, Kim and Choi, 2015), with the most common stress-related disorders being posttraumatic stress disorder (PTSD), anxiety disorders and major depressive disorder (MDD) (Smoller, 2016). Despite the established importance of stress in disease, we currently lack objective measures of the biological impact of stress on the body. A better understanding of the metabolic changes caused by stress is crucial for elucidating the mechanisms underlying the development of stress-related disease and developing better treatments (Lee, Kim and Choi, 2015). Furthermore, personalised medicine requires biomarkers that are sufficiently sensitive and specific for diagnosing potentially damaging stress exposure and predicting future outcomes (Dhama *et al*., 2019).

Elevated cortisol, which can be measured non-invasively in saliva or hair, is one of the most common measures currently used to assess acute and chronic stress. However, due to the inverted U-shaped relationship between cortisol levels and numerous neurobiological and behavioural endpoints, cortisol is nonspecific and cannot be used to reliably distinguish understimulation from overstimulation and stress (Sapolsky, 2015). Moreover, there are individual differences in the level of stress associated with optimum stimulation for health and well-being with individuals vulnerable to stress-related disorders having lower optimum cortisol levels than individuals resilient to stress (Sapolsky, 2015).

Due to the complexity of the stress response, a more holistic approach is required, moving away from the measurement of single biomarkers such as cortisol. Triangulation, whereby multiple biomarkers, each with its own strengths and weaknesses, are measured simultaneously, should deliver a more reliable estimate of an individual’s condition. Untargeted metabolomics identifies and compares the relative concentrations of a large cohort of metabolites (small molecules < 1,500 Da) between samples of interest, yielding a specific metabolic signature or fingerprint of a given exposure or manipulation (Goodacre *et al*., 2004; Wishart, 2008; Chen, Zhong and Zhu, 2020). The results can be used to identify a panel of metabolites that effectively discriminate different conditions. We test the hypothesis that untargeted metabolomics conducted on samples obtained before and after exposure to a major stressor will reveal new biomarkers and shed light on the metabolic pathways involved in the etiology of stress-related diseases.

Collecting samples longitudinally before and after a terrifying, life-threatening event, characteristic of a major stressor, is practically and ethically challenging. Here we make use of an opportunistic pig model that we argue has strong ecological validity for the types of events known to potentially result in PTSD and other stress related disorders. Handling of pigs pre-slaughter involves a 24-hour period during which animals are transported, re-mixed with unfamiliar pen-mates and held in a new environment where they are exposed to the noise and smell of the slaughterhouse (Rubio-González *et al*., 2015). These events result in, prolonged thirst and hunger, restricted movement, negative social behavior, resting problems, fatigue, pain, fear and respiratory distress (Nielsen *et al*., 2020), with physiological consequences sufficient to impair meat quality (Rubio-González *et al*., 2015; He *et al*., 2019). Pigs have physiological similarities to humans making them a common animal model in biomedical research (Gieling, Nordquist and van der Staay, 2011; Henze *et al*., 2019). Additional benefits of the model include the homogeneous genetic background, age, nutrition and rearing environment of the animals, all of which reduce random variation in physiology. Saliva samples were collected non-invasively from standard environmental enrichments provided in the animals’ pens, ensuring that baseline samples were uncontaminated by sampling stress.

## RESULTS

### Effects of 24-hours of stress on the saliva metabolome

For the examination of acute stress as detailed in Figure 1, longitudinal saliva samples were collected non-invasively from 200 pigs by providing cotton ropes for the animals to chew as environmental enrichment (Morgan *et al*., 2018, 2019; Morgan and Raz, 2019). The pigs were penned in groups and each rope provided a pooled sample from 5-10 animals. Saliva samples were collected from the pigs twice: (1) 24 hours before slaughter, while pigs were still in their pen, in a familiar group from weaning (n=31 samples); and (2) after short transport, regrouping with unfamiliar pen-mates, and 24 hours holding in the new environment of the slaughterhouse (n=32 samples). All samples were kept on ice, and stored at -80°C within two hours of collection until the metabolomics analyses. An untargeted metabolomics approach was used involving ultra-high-performance liquid chromatography coupled with high-resolution mass spectrometry (UHPLC-HRMS). Metabolites were extracted from the saliva samples using an extraction procedure (methanol:acetonitrile) suitable for polar metabolites. UHPLC-HRMS was performed using a hydrophilic interaction liquid chromatography (HILIC) analytical column, in both positive and negative ionization mode for broader metabolite coverage. A pooled sample from all samples was injected from the same vial for quality control (QC), as well as pooled samples extracted four times for quality assurance. Overall, obtained UHPLC-HRMS data revealed the relative quantification of 2,518 metabolite features in total, in the positive ionization mode, and 1,165 metabolite features in the negative ionization mode. After data evaluation and cleaning, including background spectral filtering (noise elimination) and QC based filtering (metabolite features with coefficient of variation (CV) ≥ 30% were removed), 1,564 metabolite features from the positive mode and 850 metabolite features from the negative mode remained for further statistical analyses. Of these, 920 metabolite features were putatively annotated in the positive ionization mode, and 531 in the negative ionization mode.

**Fig. 1.**
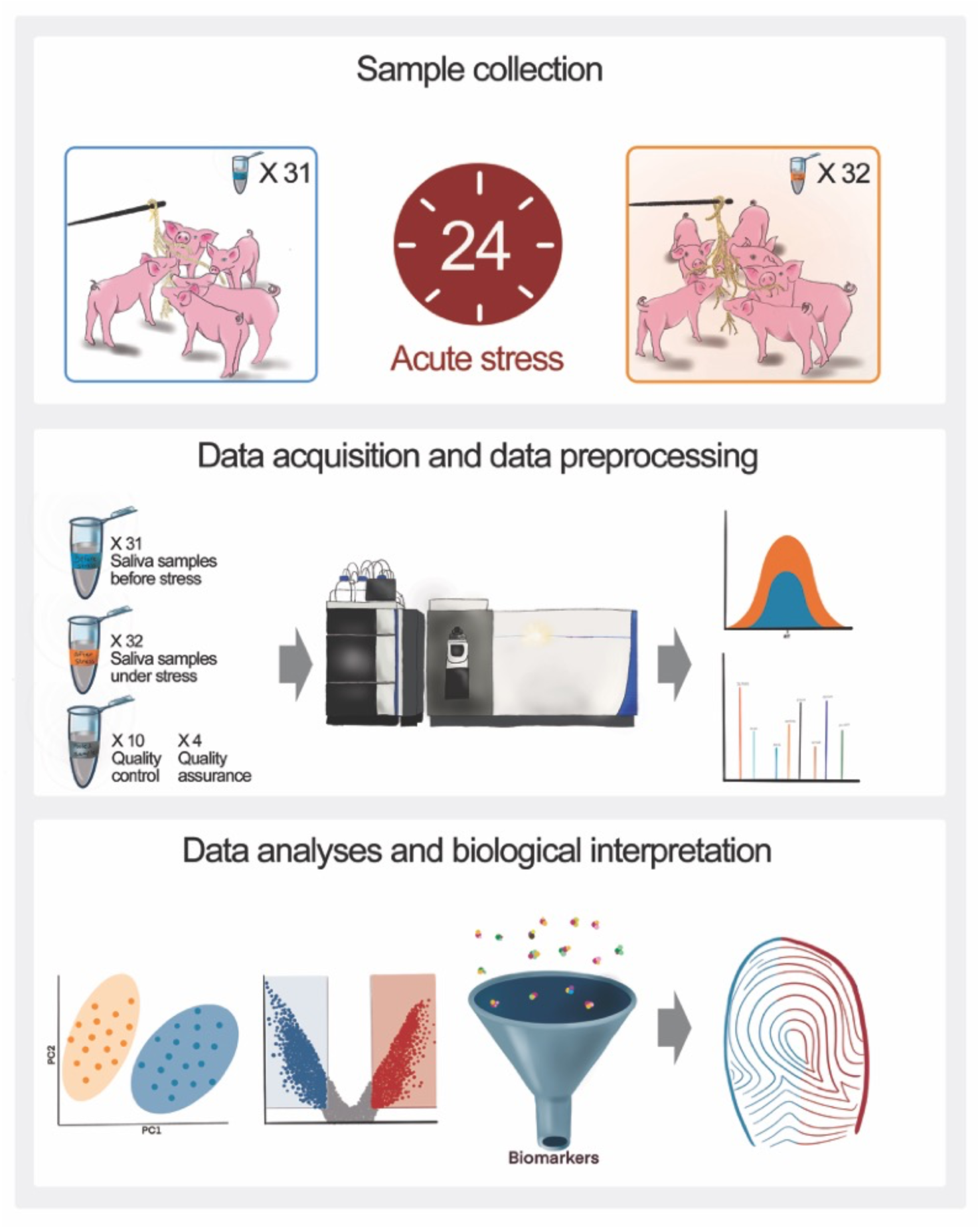
Study design. For the examination of acute stress, saliva samples were obtained non-invasively from 200 pigs. Cotton ropes were provided, yielding 63 pooled saliva samples, each from 5-10 pigs. Saliva was collected from the same pigs twice: (1) before stressful conditions (n=31) ; and (2) after 24 hours in stressful conditions (n=32). Metabolites were extracted and analysed using an ultra-high-performance liquid chromatography high-resolution mass spectrometer. Raw data was used for data pre-processing, identification of features as metabolites, and data analyses for the identification of altered metabolites and metabolic pathways.

A principal component analysis (PCA) score plot revealed clear separation between pig saliva metabolites before and after the acute stress intervention, suggesting a clear influence of the intervention on the saliva metabolome (Fig. 2).

**Fig. 2.**
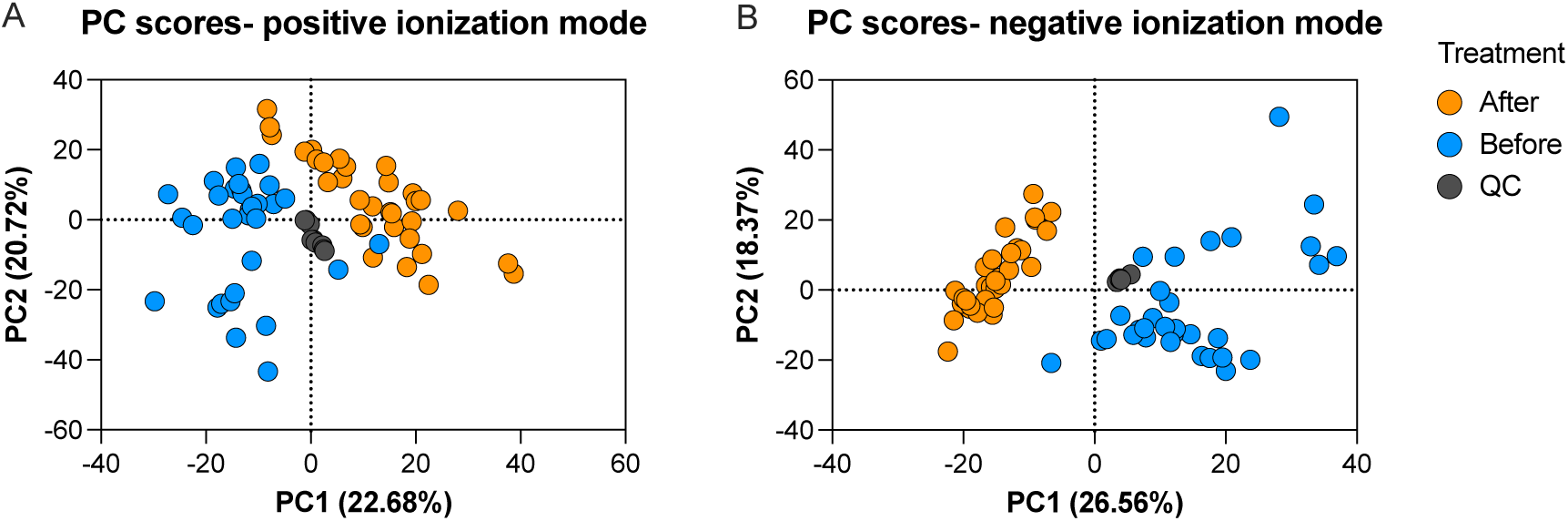
PCA plot of pig saliva metabolome. A principal component analysis (PCA) plot of the metabolome data that characterizes pig saliva before (Blue) and after (Orange) acute stress. Each dot represents a sample and the colour represents the type of sample. Repeatability (grey) is demonstrated by pooled samples, injected along the analyses as quality controls. **(A)** A PCA score plot of metabolite features from data acquisition in positive ionization mode. **(B)** A PCA score plot of metabolite features from data acquisition in negative ionization mode.

Furthermore, relative concentrations of 932 metabolite features in the positive ionization mode and 591 metabolite features in the negative ionization mode were significantly different before and after the stressful intervention (Fig. 3; *Adj* P < 0.05). Of these features, those with at least a two-fold change in relative concentration were considered differentially affected by stress. Accordingly, in the positive ionization mode 243 metabolite features were significantly upregulated (a fold change of 2.00 to 17.12) after stress, and 357 metabolite features were significantly downregulated (a fold change of 2.0 to 333.33) after stress (Fig. 3A; *Adj* P < 0.05). In the negative ionization mode, 49 metabolite features were significantly upregulated (a fold change of 2.01 to 53.05) after stress, and 333 metabolite features were significantly downregulated (a fold change of 2.04 to 250.0) under stress (Fig. 3B; *Adj* P < 0.05).

**Fig. 3.**
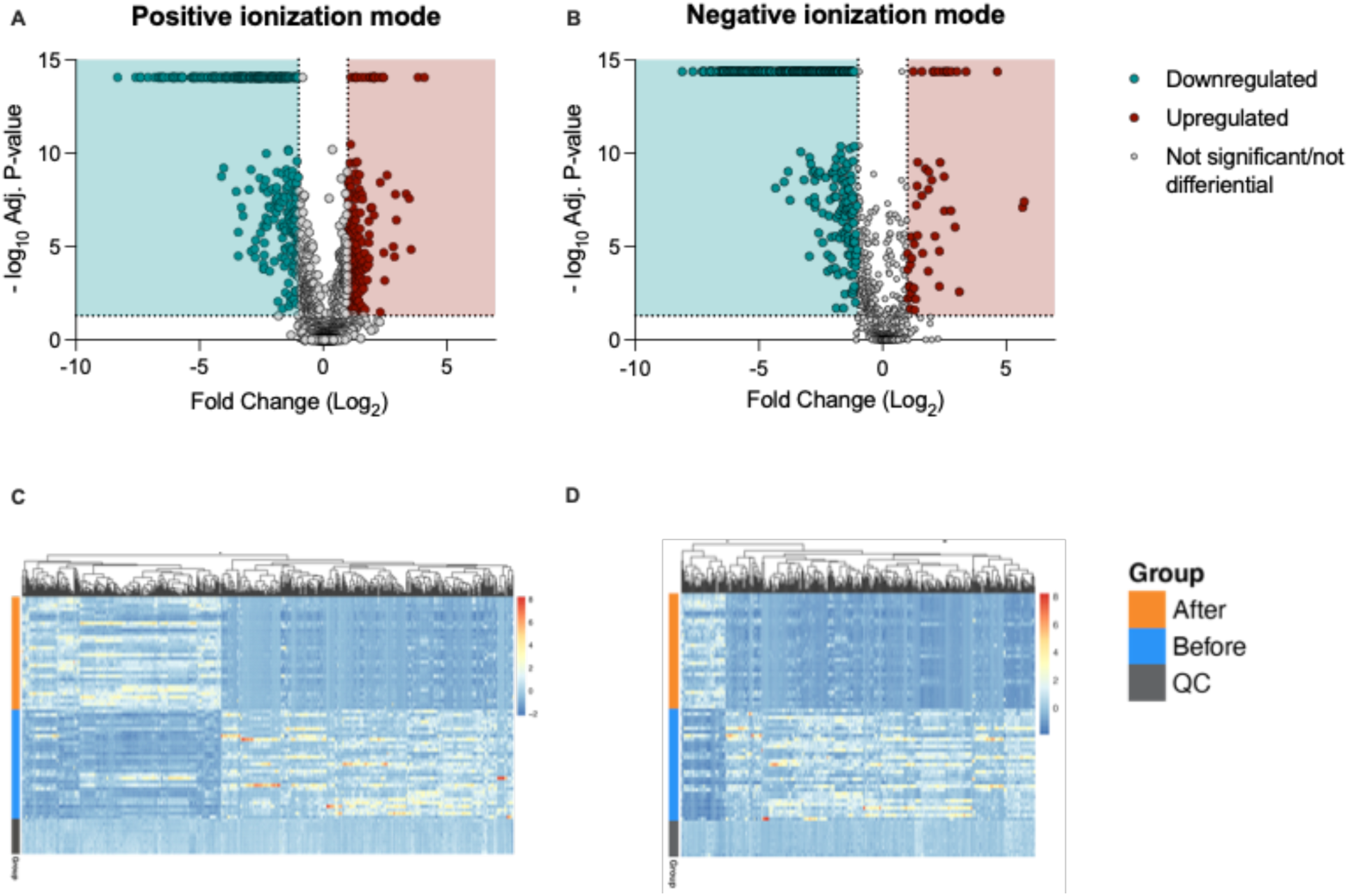
Differential metabolites features after stress. **(A-B)** Differential analysis of metabolite ratios between pigs in their familiar environment (Before) and 24 hours after (After) the onset of stress. **(A)** Volcano plot of metabolite features from data acquisition in positive ionization mode. **(B)** Volcano plot of metabolite features from data acquisition in negative ionization mode. Each dot represents a metabolite feature. Significant upregulated metabolite features are in red on the right (2 < fold change; *Adj* P < 0.05). Significant downregulated metabolite features are in blue on the left (fold change < 0.5; *Adj* P < 0.05). **(C-D)** Heat map analysis of differential metabolite features (fold change < 0.5; 2 < fold change; *Adj* P < 0.05), **(C)** in both positive and **(D)** negative ionization modes. Each row represents a sample and each column a metabolite feature. Purple indicates that metabolite features were downregulated, and yellow indicates that metabolite features were upregulated.

Follow-up analyses included investigation of metabolite features responsible for the clustering of the metabolome changes before and after the stressful intervention. The complete screening for biomarkers as summarized in Table 1, and fully detailed in Table S1. As mentioned, we searched for metabolite features for which the relative concentrations after stress were not only significantly different, but also showed at least a two-fold change (*Adj* P < 0.05; 2 ≤ FC or FC ≤ 0.5). Each of these metabolite features that could serve as potential biomarkers was manually checked for the match of the precursor ion mass (*m/z*) and fragments (MS/MS) to online and in-house libraries. Metabolite identification confidence level for each identified metabolite in this study was assigned following the modified four-level classification scheme from the Metabolomics Standards Initiative (Sumner *et al*., 2007; Salek *et al*., 2013): level 1- match to commercial standards; level 2- full match to online databases of the precursor ion and its fragments (both *m/z* and MS/MS); level 3- putative annotation based on precursor ion; and level 4- not annotated). To ensure higher confidence, the metabolites discussed as potential biomarkers in this paper are only those with identification confidence levels 1 and 2. These, were mainly decreased amino acids and metabolites synthesized endogenously by amino acids and B vitamins. In addition, the most promising biomarkers that were not identified with high confidence, or were not annotated at all, are detailed in Table S1.

**Table 1.** Summary of potential biomarkers. Summary of significantly differential metabolite features after acute stress. Metabolites with at least a two-fold change ratio before and after stress were considered differential and *Adj* P < 0.05. Metabolite features are divided according to their fold-change ratios differences, as well as the assigned level of confidence in the annotation (level 1- match to commercial standards, level 2- full match of the precursor ion and its fragments to online databases (both *m/z* and MS/MS), level 3- putative annotation based on precursor ion (*m/z*), level 4- not annotated). Further detail is provided in the supplementary information Table S1.

|  |  | **Total number of potential biomarkers** | Level 1 | Level 2 | Level 3 | Level 4 |
| --- | --- | --- | --- | --- | --- | --- |
| **Positive ionization mode** |  |  |  |  |  |  |
| **Upregulated**<br>**248** | $2 \leq FC < 5$ | **234** | 0 | 5 | 66 | 163 |
| | $5 \leq FC \leq 10$ | **9** | 0 | 1 | 4 | 4 |
| | $10 < FC$ | **5** | 0 | 0 | 3 | 2 |
| **Downregulated**<br>**359** | $0.2 < FC \leq 0.5$ | **188** | 5 | 11 | 112 | 60 |
| | $0.1 \leq FC \leq 0.2$ | **82** | 2 | 2 | 53 | 23 |
| | $FC < 0.1$ | **89** | 0 | 2 | 62 | 27 |
| **Negative ionization mode** |  |  |  |  |  |  |
| **Upregulated**<br>**50** | $2 \leq FC < 5$ | **35** | 0 | 1 | 23 | 11 |
| | $5 \leq FC \leq 10$ | **11** | 0 | 0 | 4 | 7 |
| | $10 < FC$ | **4** | 0 | 0 | 1 | 3 |
| **Downregulated**<br>**333** | $0.2 < FC \leq 0.5$ | **144** | 0 | 5 | 97 | 42 |
| | $0.1 \leq FC \leq 0.2$ | **56** | 0 | 2 | 42 | 12 |
| | $FC < 0.1$ | **133** | 0 | 3 | 81 | 49 |

Another approach in the investigation of novel biomarkers and metabolic fingerprints, is to look at the alteration of entire metabolic pathways, rather than single potential biomarkers. Therefore, 144 metabolite features from both the positive and negative ionization with higher annotation confidence (level 1 and level 2; when both precursor ion and its fragments, MS/MS, were matched to in-house library of commercial standards or online databases). 95 of these putatively annotated metabolite features were successfully matched to a metabolic pathway by Metaboanalyst software 5.0 (Pang *et al*., 2021). Several metabolic pathway alterations were induced following the 24-hour stressful intervention. These pathways are represented by both -log_10_ (*Adj* P-value) > 1.1220, which is equivalent to a *Adj* P-value < 0.05 and high pathway impact score, calculated by overrepresentation of metabolites from the same pathway, as well as highly important metabolites to the pathway. Accordingly, as shown in Figure 4, histidine metabolism, phenylalanine, tyrosine and tryptophan biosynthesis, arginine and proline metabolism and arginine biosynthesis are both significantly different, as well as have relatively high pathway impact. Moreover, the pathways of nicotinate and nicotinamide metabolism, lysine degradation, alanine aspartate and glutamate metabolism and vitamin B6 metabolism were also highlighted and show promising direction for further investigation.

**Fig. 4.**
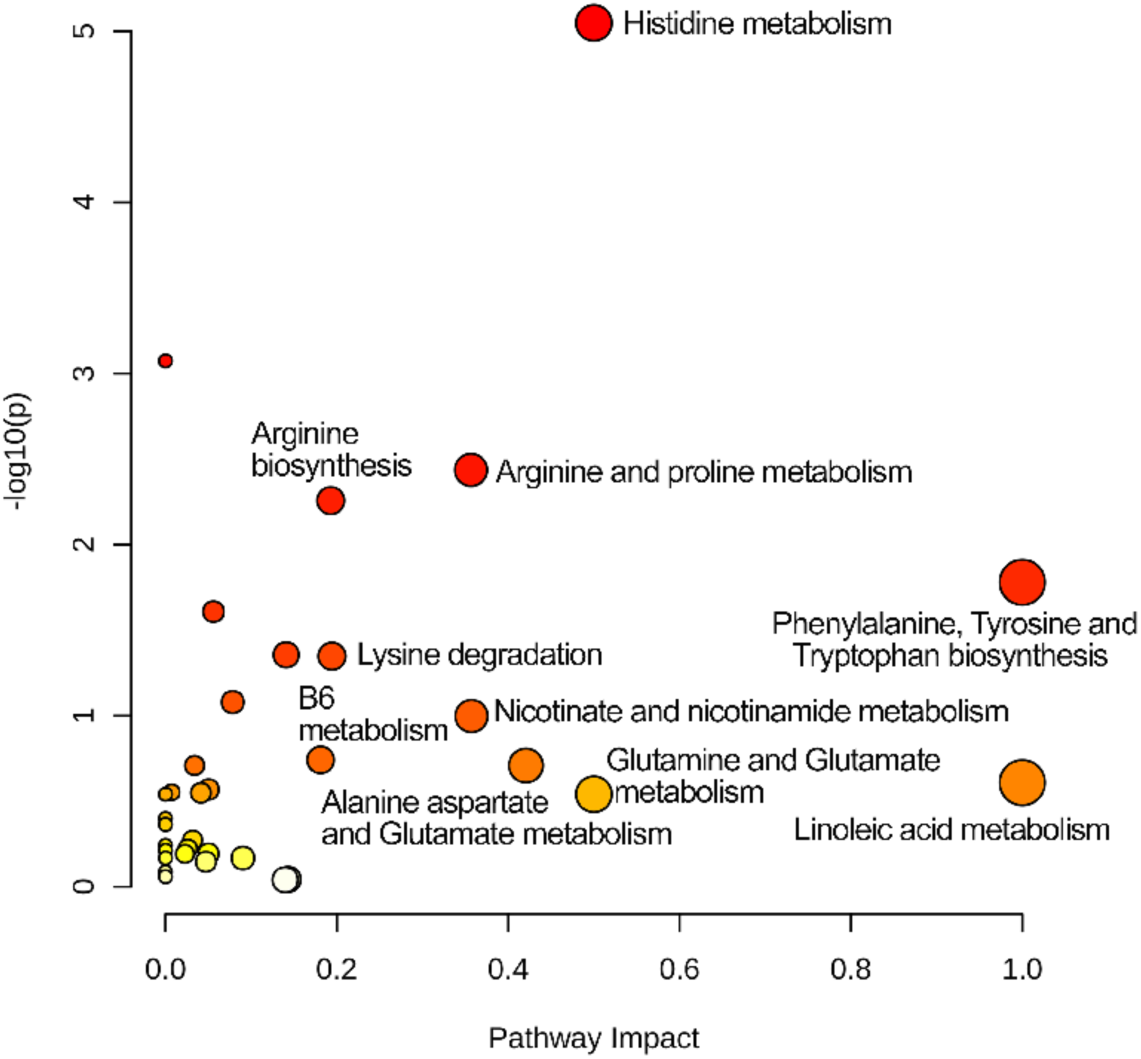
Altered metabolic pathways after acute stress. Pathway analysis plot. Dots represent metabolic pathways altered after 24 hours of major stress. The Y-axis represents the logarithm transformed P-value adjusted for multiple comparisons. The X-axis represents the pathway impact, calculated by overrepresentation of metabolites from the same pathway, as well as highly important metabolites to the pathway. The pathways that were most associated with stress are a combination of high pathway impact and -log_10_ (*Adj* P-value) > 1.1220, which is equivalent to a P-value < 0.05 (the top right corner of the graph). Colour represents significance, from white (not significant) to red (significant), and larger dot size represents a higher impact score.

## DISCUSSION

The trajectory of stress and related disorders has been widely investigated (Schneiderman, Ironson and Siegel, 2005; Lee, Kim and Choi, 2015; Yaribeygi *et al*., 2017), but there remains a gap in our understanding of the acute systemic effects of major stress (Greef, Stroobant and Heijden, 2004; Quinones and Kaddurah-Daouk, 2009; Lee, Kim and Choi, 2015). Our aim was to characterize the metabolic fingerprint of major stress, in order to provide a comprehensive understanding of its acute physiological effects and to reveal novel potential biomarkers. We utilized an untargeted metabolomics approach in which we used UHPLC-HRMS to characterize the change in the saliva metabolome of the same group of pigs measured before and 24-hours after exposure to major stress involving transport, social mixing and holding in the novel and frightening environment of a slaughterhouse. Our results provide a basis for further investigation of the impacts of stress and the mechanisms underlying the development of stress-related disorders such as PTSD in humans. Moreover, they may shed light on potential drugs and nutritional supplements for treating stress-related disorders.

In this study, we found several hundreds of metabolite features that display a 2.00 to 333.33 fold change after major stress (600 metabolite features in the positive ionization mode and 382 metabolite features in the negative ionization mode). These metabolite features (detailed in Table 1 and Table S1) are divided into two main groups; The first, are metabolite features for which the retention time, as measured by the liquid chromatography, and the molecular mass of the molecule and its fragments, as measured by the mass spectrometer, could be used to identify the feature as a specific metabolite (identification confidence levels 1 and 2). The second group, are metabolite features that are part of the metabolic fingerprint of stress and may provide potential biomarkers, but for which identification requires additional analytical methods (confidence levels 3 and 4). The main metabolic changes identified were decreases in amino acids and metabolites synthesized endogenously by amino acids and B vitamins. Many types of stress including psychosocial stress, injuries, surgeries, infection and cancer activate metabolic pathways that consume amino acids (Dadmarz *et al*., 1998; Reeds and Jahoor, 2001; C, I and D, 2002). In this study, the ratio of relative concentrations of arginine after versus before stress was 0.154, indicating that arginine was significantly lower after 24 hours of stress. Arginine is a conditionally essential amino acid, because while it can be synthesized, it becomes essential during trauma and disease (Reeds and Jahoor, 2001; Dasgupta, Hebbel and Kaul, 2006; Huynh and Chin-Dusting, 2006). Lysine and tryptophan also decreased after stress in this study (ratios of 0.162 and 0.492 respectively). Prolonged lysine insufficient diets are correlated with impaired quality of life and mental health (Smriga *et al*., 2004). Thus, a decrease under prolonged stress might be consequential, especially if the diet is deficient in lysine. Tryptophan is also an essential amino acid, which functions as a precursor of serotonin, as well as in the synthesis of kynurenine. The tryptophan-kynurenine pathway is associated with cognitive symptoms and neurodegenerative diseases (Jenkins *et al*., 2016). While the role of serotonin in regulating mood and cognition is known, the complete mechanism of kynurenine and the unwanted symptoms of neurodegenerative diseases is not fully discovered (Jenkins *et al*., 2016). There is controversial evidence in the literature, regarding the influence of tryptophan and its supplementation on diet on mood change and stress related disorders such as major depression (Merens, Willem Van der Does and Spinhoven, 2007; Soh and Walter, 2011). B vitamins also decreased in our study; the ratio of relative concentrations of nicotinamide after stress versus before was 0.498. Nicotinamide, the amide form of vitamin B3, has a role as a precursor in the synthesis of coenzymes nicotinamide adenine dinucleotide (NAD+) and nicotinamide adenine dinucleotide phosphate (NADPH), which are responsible for various functions including oxidative deamination and lipid catabolism (Oblong, 2014). Nicotinamide has a key role in development, growth and maintenance of the central nervous system and is also considered to have protective effects on the nervous system, providing protection from neurodegenerative diseases such as Alzheimer’s, Parkinson’s and age-related macular degeneration (AMD) (Fricker *et al*., 2018). Another B vitamin that was decreased in this study after stress was pyridoxine, vitamin B6 (0.359). Pyridoxine has a key role in amino acids metabolism and cell functioning (Parra, Stahl and Hellmann, 2018). Moreover, as for other B vitamins, pyridoxine is considered to have protective effects against cognitive decline and the related pathologies such as Alzheimer’s disease (Smith and Refsum, 2016). The decrease in these three metabolites (tryptophan, nicotinamide and pyridoxine) under acute stress may be important in elucidating the effect of stress on the later trajectories. Additional metabolites that may shed the light on these trajectories, were also decreased under stress in this study. Carnosine decreased after stress (ratio of 0.467). Carnosine, a dipeptide, which is endogenously synthesized from the amino acids alanine and histidine, has many physiological functions and is dominant in tissues such as the brain, heart, muscles, liver, brain, and kidneys (Wu *et al*., 2013). L-carnosine has been shown to function as an antioxidant, an anti-inflammatory, an accelerator of wound healing and it might even have antidepressant effects (Bellia, Vecchio and Rizzarelli, 2011). Furthermore, L-carnosine has been shown to act as an inhibitor of the toxic amyloid-β peptide (Aβ) present in the vascular plaques in Alzheimer’s patients, as well as in reducing aging-related mitochondrial dysfunctions (Preston *et al*., 1998; Bellia, Vecchio and Rizzarelli, 2011; Corona *et al*., 2011; Wu *et al*., 2013). Hypoxanthine which also decreased under stress (0.416) is a derivative of purines, and is associated with abnormal purine metabolism in Alzheimer’s patients (Liang *et al*., 2015). Agmatine, which is also induced endogenously as part of the stress and inflammatory response, has a role as a neuromodulator. It has been suggested that agmatine has a protective effect in the development of trauma and stress related disorders, such as PTSD, anxiety, and depression (Halaris and Plietz, 2012). Moreover, it may have protective effect against neurodegenerative diseases such as Alzheimer’s and Parkinson (Gilad *et al*., no date; Song *et al*., 2014, 2016). In this study after 24 hours in stressful conditions, saliva agmatine also decreased (ratio of 0.377). It has been shown that as a result of stress, trauma and inflammation, agmatine level is increased, but cannot compensate fully for its harmful impact, at least not the endogenous production (Zhu *et al*., 2008; Gawali *et al*., 2017). We hypothesize that the agmatine that was produced under stress was consumed as part of the protective mechanism for neural functioning, and therefore was already decreased after 24 hours in stressful conditions. It has been described that agmatine is elevated after the first few hours from stressful exposure, but the exact mechanism of agmatine as part of the stress response, as well as the longer term effect remained unclear (Aricioglu, Regunathan and Piletz, 2003). Moreover, it was also demonstrated in rodents that agmatine has anxiolytic effect, as well as suppression of depression-like behaviour (Gong *et al*., 2006; Bahremand *et al*., 2018). Further investigation requires measuring the levels in saliva and other biofluids, as well as in the brain, from the stressful exposure, and after different time periods, in order to get a comprehensive understanding.

The main limitation of this study, as in all untargeted metabolomics studies, is the failure to identify several metabolite features. Identification of metabolites is the main bottleneck in the workflow in untargeted metabolomics. Identification is based on the mass of the intact metabolite (more specifically, the mass to charge ratio, *m/z*), its fragments (MS/MS), and the retention time in the analytical column. Although we can record this information for thousands of metabolite features in one analysis, still many of the metabolite features remain unidentified as a specific metabolite. This limitation is due to the lack of available standards, limited databases and the available algorithms (Domingo-Almenara *et al*., 2018). It is likely that several unidentified metabolites have the potential to serve as novel biomarkers and are thus worthy of further investigation. For example, Table S1 includes metabolite features that are significantly differentially regulated after stress, with at least a ten-fold change ratio (*Adj* P < 0.05; FC > 10; FC < 0.1). These features are putatively annotated as level 3 or not annotated at all (level 4). One future solution to the large number of unidentified metabolite features will be a cross-species analysis of the effects of stress. Triangulating the overlap in the metabolite features altered by different types of stress, in different species, and in different biological matrices, might help to narrow down the list of potential biomarkers from hundreds, to a shorter list of interest for further identification and validation.

In light of all the above, this research shed light on the metabolic changes under acute stress, by metabolomics analyses in a unique pig saliva model. It lays foundations for further research for the identification of the unidentified metabolite features, as well as provide a new perspective about stress. Previous studies have focused on upregulated metabolites, such as cortisol, that are elevated under stressful conditions. The current research emphasizes a decrease in several metabolites that are downregulated under acute stress. The decrease in these metabolites, several of which might have protective effects on the nervous system, reducing the likelihood of neurodegenerative diseases and psychological disorders, suggests a novel future direction in research on trauma and stress-related disorders. Perhaps, focusing on downregulated metabolites, as we did in this research, would lead to new future discoveries. This is similar to what has been suggested regarding stress-related disorders, whereby focusing on the mechanism of resiliency, rather than the stress, may bridge this gap of knowledge and lead to new therapeutic avenues (Russo *et al*., 2012; Kalisch *et al*., 2017). Clinically, focusing on these downregulated metabolites as prophylactic treatment may allow individuals to cope better with stress, and stay for longer in the left side of the inverted U-shape, as resilient, before getting to the turning point of vulnerability.

In summary, our study indicates that after acute stress of 24 hours, in a unique pig model, while several metabolites are upregulated, a higher number of metabolites are downregulated, and their related metabolic pathways changed accordingly. The main metabolic changes that we identified were a decrease in amino acids and metabolites synthesized endogenously by amino acids and B vitamins, as well potential unidentified metabolite features that require further investigation. The metabolic fingerprint of stress that we reveal provides new insights into the trajectory of stress and the related disorders. This research demonstrates a clear future direction for the investigation of stress and resilience, as well as potential therapeutics and nutritional targets.

## MATERIALS AND METHODS

### Sample collection

Saliva samples were collected non-invasively from 200 pigs, ages ∼165 days, mixed breed of Landrace, Large-White, Pietrain and Duroc. The study was performed at Lahav Animal Research Institute (LAHAV C.R.O; Kibbutz Lahav, Israel), according to the Tel Aviv University Institutional Animal Care and Use Committee. The pigs were housed in groups and each saliva sample represented a pooled sample from 5-10 pigs. Saliva was collected by providing 100% cotton ropes as environmental enrichment, similar to the method recently published (Morgan *et al*., 2018, 2019; Morgan and Raz, 2019). Saliva was collected from the same 200 pigs twice: (1) 24 hours before slaughter, while pigs were still in their pens, in their familiar environment (n=31). (2) After a short transport to the slaughterhouse (10 minutes including loading time), regrouping with unfamiliar pen-mates, and 24 hours wait in new environment at the slaughterhouse (n=32). Although the same 200 pigs are represented in the samples collected both before and after stress, the individual samples represent different combinations of individuals before and after stress due to the regrouping that occurred. Thus, the samples represent independent pooled samples from the same population of pigs collected before and after stress. Saliva samples were stored immediately after collection on ice, and within two hours aliquoted and stored at -80°C until the extraction procedure and data acquisition.

### Sample preparation

Metabolites were extracted from the pig saliva samples using an extraction procedure targeting primarily polar metabolites, similarly to recently published methods (Palmer, Cooper and Dunn, 2019; Southam *et al*., 2020). Saliva samples were thawed on ice and briefly vortexed (10 seconds). Saliva samples were centrifuged at 1300 rpm, 4°C, for 10 minutes and the clear fluids were transferred to new 1.5 mL laboratory centrifuge tubes. Samples were vortexed again and 50 μL of each sample, including six QC samples were transferred to a new 1.5 mL laboratory centrifuge tube (from a pooled sampled prepared from 75 µL from each sample in a 15 ml tube as a pooled QC vial. Then, 150 µL of ice-cold 50:50 methanol:acetonitrile (v/v) was added to each. Next, vortex for 15 seconds and centrifugation at 14,000 rpm, at 4°C for 20 minutes was performed. Finally, 150 μL from each sample were transferred to LC vials with inserts for immediate analysis. Samples were injected in randomized order with quality control samples every eight injections. Injection volume was 5 µL, and autosampler compartment was kept at 7°C.

### Quality assurance and quality control

In order to assure the quality of the data, several actions took part: 1) A pooled quality control sample (QC) was prepared by taking 75 µL from each experimental sample, injected from the same vial every eight samples. The pooled QC sample injected along the sequence, serves as quality control for the instrument performance. 2) A pooled QC sample was extracted four times (QA), injected randomly along the sequence as quality assurance for the extraction procedure. 3) Extracted water sample was used as a blank, to subtract background ions in data analysis. R package RawHumus (Dong *et al*., 2022) was used to evaluate the data quality based on QC samples. The resulting QC reports are shown in File S2 for QC data obtained in positive ionization mode and File S3 for QC data obtained in negative ionization mode.

### Ultra-high-performance liquid chromatography-tandem mass spectrometry

High performance liquid chromatography was performed by Vanquish UHPLC using analytical Accucore™ 150 Amide HILIC column (100 × 2.1 mm, 2.6μm), coupled to a high-resolution mass spectrometer Orbitrap Q-Exactive™ Focus (Thermo Scientific™) equipped with a HESI ion source. All samples were first analysed in positive ionization mode and then injected again for analysis in the negative ionization mode. Mass calibration was performed on the day of starting the data acquisition.

Positive ionization mode: mobile phase A: 10 mM ammonium formate, 0.1% formic acid, 95% acetonitrile/water (v/v). Mobile phase B: 10 mM ammonium formate, 0.1% formic acid, 50% acetonitrile/water (v/v). Gradient: t= 0.0min, 1% B, t= 1.0min, 1.0% B, t=3.0min, 15.0% B, t=6.0min, 50.0% B, t=9.0min, 95% B, t=10.0min, 95% B, t=10.5min, 1% B, t=15.0min, 1% B Negative ionization mode: mobile phase A: 10 mM ammonium acetate, 0.1% acetic acid, 95% acetonitrile/water (v/v). Mobile phase B: 10 mM ammonium acetate, 0.1% acetic acid, 50% acetonitrile/water (v/v). Gradient: t= 0.0min, 1% B, t= 1.0min, 1.0% B, t=3.0min, 15.0% B, t=6.0min, 50.0% B, t=9.0min, 95% B, t=10.0min, 95% B, t=10.5min, 1% B, t=15.0min, 1% B. For both ionization modes, the column temperature was 35 °C and injection volume was 5 μL.

Tune file settings: Sheath gas flow rate 30 (arbitrary units), aux gas flow rate 13 (arbitrary units), spray voltage 3.2kV (2.7kV negative ionization mode), capillary temperature 350° C, aux gas heater temperature 400° C.

### Mass spectrometric settings

Full scan mass resolution 35,000, Scan range 70 to 1050 m/z, ACG target 1e^6^, spectrum data type: profile. dd-MS2 discovery mode, MS2 mass resolution 17,500, isolation window 3.0 Da, collision energy 30 eV.

### MS-data preprocessing and data analysis

Post-acquisition data processing was accomplished using Compound Discoverer™ (Thermo Scientific™) software 3.2, commonly used for untargeted metabolomics analyses. In brief, raw data was uploaded and processed; missing values imputation was automatically performed by the software. Then, features with more than 20% missing values across sampled were filtered. The obtained mass to charge ratio (*m/z)*, including MS/MS fragments and retention time were searched against large spectral libraries for metabolite annotation. After data evaluation and cleaning process, including background spectral filtering (noise elimination) and quality control (QC)-based filtering (Metabolite features with CV ≥ 30% were removed), the dataset was suitable for statistical analysis. First, Principal Component Analysis (PCA), as an unsupervised statistical method, was utilized. Then, the screening for biomarkers was done by step-wise filtering of *Adj* P-value <0.05, and different fold change ratios as detailed in the results section. Data exploration also included further investigation of the affected metabolic pathways by implemented KEGG Pathway Database and Metabolika pathway analysis into Compound Discoverer™, as well by the online tool of MetaboAnalyst 5.0 software (Pang *et al*., 2021). Finally, the detected potential biomarkers were confirmed manually by spectra evaluation and comparison to the fragmentation pattern and retention time of commercial standards or online library mzCloud™ and commercial standards by Sigma. Adjusted P-value < 0.05 was reported as significant.

## Supplementary materials

File S1. The complete list of significantly different metabolite features

File S2. Evaluation of QC data obtained at positive ion mode

File S3. Evaluation of QC data obtained in negative ion mode

## ACKNOWLEDGMENTS

We thank Lahav CRO for allowing us to conduct the research on their farm, as well as to the whole farms’ team.

## Author contributions

Conceptualization: LM, MB, EG, RB, SSN, YD

Methodology: LM, EG, MB, RB, YD, SSN, TW

Investigation: LM

Visualization: LM, YD

Project administration: LM

Supervision: MB, EG, JZ, HC, BY, LAA

Writing – original draft: LM

Writing – review & editing: LM, RB, SSN, LC, TW, YD, BY, LAA, HC, JZ, MB, EG

## Data availability

All data analysed during this study is included in this paper and its Supplementary files.

