## Supplementary material for "Saliva metabolome alterations after acute stress": File S2: QC_Report_pos (1).html

RawHumms Data Quality Control Report


### RawHumms Data Quality Control Report

###### 14, February, 2022

### 1 Introduction

Robust and reproducible data is essential to ensure confidence in analytical results and is particularly important for large-scale metabolomics studies. Therefore raw data need to be inspected before data processing and statistical analysis in order to detect measurement bias and verify system consistency. In liquid chromatography mass spectrometry (LCMS) based metabolomics studies, proper quality control (QC) checks are particularly important to ensure reliable and comparable results within experimental measurements [1-2].

**RawHummus** is an user-friendly web application for rapid data quality check based on raw QC samples. It generates an HTML report with interactive plots, statistics and illustrations that help users evaluate their data quality and LCMS system performance.

### 2 Chromatogram

#### 2.1 TIC plot

`Total Ion Current (TIC)` chromatogram represents the summed ion intensity across each scan in the analysis. **The interactive overlaid TIC plot** can be used for rapid inspection of retention time (RT) and ion intensity fluctuations.

#### 2.2 Summed TIC Plot

**Summed TIC plot** is another quick-and-dirty way to overview global ion intensity variations among QC samples. It summed TIC across the entire points (scans) from the analysis. **Dashed red line** is **mean** of summed TIC and **blue lines** represent **mean + 2SD** and **mean - 2SD**, respectively.

#### 2.3 TIC Correlation Analysis

`Pearson correlation` is used to quantify the metabolic profile similarity among QC samples. Pearson correlation coefficient (**R**) over **0.85** indicate high metabolic profile similarity in RT and chromatogram peak shape. If the value is below **0.85**, it will be highlight in **red** in Table 1.

> **Note that** RT were binned by 0.1 min for Pearson correlation calculation.

Table 1: Metabolic profile similarity

|  | QC\_p..1..mzML | QC\_p..10..mzML | QC\_p..2..mzML | QC\_p..3..mzML | QC\_p..4..mzML | QC\_p..5..mzML | QC\_p..6..mzML | QC\_p..7..mzML | QC\_p..8..mzML | QC\_p..9..mzML |
| --- | --- | --- | --- | --- | --- | --- | --- | --- | --- | --- |
| QC\_p..1..mzML | 1 | 0.992 | 0.998 | 0.999 | 0.999 | 0.998 | 0.997 | 0.984 | 0.991 | 0.992 |
| QC\_p..10..mzML | 0.992 | 1 | 0.989 | 0.991 | 0.989 | 0.989 | 0.989 | 0.992 | 1 | 0.999 |
| QC\_p..2..mzML | 0.998 | 0.989 | 1 | 0.998 | 0.998 | 0.997 | 0.996 | 0.981 | 0.988 | 0.989 |
| QC\_p..3..mzML | 0.999 | 0.991 | 0.998 | 1 | 0.999 | 0.998 | 0.998 | 0.983 | 0.99 | 0.991 |
| QC\_p..4..mzML | 0.999 | 0.989 | 0.998 | 0.999 | 1 | 0.999 | 0.998 | 0.981 | 0.988 | 0.99 |
| QC\_p..5..mzML | 0.998 | 0.989 | 0.997 | 0.998 | 0.999 | 1 | 0.999 | 0.982 | 0.988 | 0.989 |
| QC\_p..6..mzML | 0.997 | 0.989 | 0.996 | 0.998 | 0.998 | 0.999 | 1 | 0.983 | 0.988 | 0.989 |
| QC\_p..7..mzML | 0.984 | 0.992 | 0.981 | 0.983 | 0.981 | 0.982 | 0.983 | 1 | 0.992 | 0.991 |
| QC\_p..8..mzML | 0.991 | 1 | 0.988 | 0.99 | 0.988 | 0.988 | 0.988 | 0.992 | 1 | 0.999 |
| QC\_p..9..mzML | 0.992 | 0.999 | 0.989 | 0.991 | 0.99 | 0.989 | 0.989 | 0.991 | 0.999 | 1 |

> **Note that** `TIC Correlation Analysis` is mainly used to evaluate peak shape similarity and RT consistency. QC files with ion intensity drift but similar profiles could still have good Pearson correlation coefficient.

# 3 MS1

#### 3.1 Auto Peaks Evaluation

In order to accurately monitor variations in mass, RT and ion intensity, **RawHummus** automatically selects **6** peaks across the entire RT range, and use them to evaluate LCMS system.

Below are the Extracted ion chromatogram (EIC) of the 6 selected ions. You can interactively view, inspect and compare them among different QC samples.

**RawHummus** performs a simple statistics to make rapid evaluation. The table below summarized the comparison result, in which maximum RT difference, mass difference and ion intensity difference, and Intensity CV are given.

`Max RT Diff (min)`: is the maximum retention time variation (in min unit). Small value indicates a good retention time consistency. If the maximum retention time variation is over **1 min**, the value will be highlight in **red** in Table 2.

`Max Mass Diff (ppm)`: is the maximum mass variation (in ppm unit). Small values indicate good mass accuracy. If the maximum mass variation is over **5 ppm**, the value will be highlight in **red** in Table 2.

`Max Intensity Fold Change`: is the maximum ion intensity variation. The value closer to 1 suggests that the ion intensity is stable. If the maximum intensity ratio is over **2**, the value will be highlight in **red** in Table 2.

If a peak is missing in some of your samples, **NA** values will be give in the table. You need to carefully inspect the peak so as to evaluate the reproducibility.

`Intensity CV (%)`: is the ion intensity coefficient of variation (also termed as relative standard deviation, STD). Smaller value indicates better ion intensity consistency. If the intensity CV is over **30%**, the value will be highlight in **red** in Table 2.

Table 2: Summary of auto-selected peaks

| Peak | Max RT Diff (min) | Max Mass Diff (ppm) | Max Intensity Ratio | Intensity CV (%) |
| --- | --- | --- | --- | --- |
| RT: 0.23 mz: 124.087 | 0.24 | 3.63 | 4.54 | 64.38 |
| RT: 2.94 mz: 114.066 | 0.06 | 1.34 | 1.06 | 2.07 |
| RT: 6.92 mz: 118.086 | 0.03 | 1.49 | 1.12 | 4.02 |
| RT: 7.5 mz: 132.077 | 0.02 | 1.27 | 1.23 | 7.46 |
| RT: 10.03 mz: 167.013 | 0.26 | 1.37 | 1.21 | 7.85 |
| RT: 14.68 mz: 83.06 | 1.2 | 1.38 | 1.66 | 18.22 |

#### 3.2 User defined peaks

Additionally, users could add their peaks of interests for inspection and comparison. If these peaks are defined in **RawHummus** and are found in the data. Similar plots and a data summary table will be given below. Otherwise, this section will be left blank.

> **Note that** noise peaks can be used to minitor the mass accuracy variation, but they may not work well to evalute RT and ion intensity variation.

```
## [1] "User defined peaks are not found"
```

# 4 MS2

MS/MS fragmentation is important for metabolite identification. RawHummus is also able to identify problems with regard to fragmentation.

> **Note that** if your data files do not contain any MS/MS information, this section will be left blank.

#### 4.1 Number of MS2 Events

`Number of triggered MS/MS`: is the total number of MS/MS events in the sample. Similar number of triggered MS/MS events indicates good reproducibility.

Table 4: Summary of MS2 events

| filename | MS2 Events |
| --- | --- |
| QC\_p (1).mzML | 4024 |
| QC\_p (10).mzML | 3957 |
| QC\_p (2).mzML | 4075 |
| QC\_p (3).mzML | 4028 |
| QC\_p (4).mzML | 4053 |
| QC\_p (5).mzML | 4034 |
| QC\_p (6).mzML | 4001 |
| QC\_p (7).mzML | 3851 |
| QC\_p (8).mzML | 3896 |
| QC\_p (9).mzML | 3889 |

#### 4.2 Precursor Distribution across Mass Range

`Precursor Distribution across Mass` plot is used to visualize the mass range proportion of fragmented precursors.

- It can be used to check peaks at which mass ranges are mainly fragmented.
- High similarity in `Precursor Distribution across Mass` among QC samples indicates good reproducibility in MS2.

```
## $`1`
```

```
## 
## $`2`
```

`Cosine Similarity` is used to quantify the similarity of precursor distribution across mass range plots among QC samples. Cosine similarity over 0.85 indicate high similarity. If the value is below 0.85, it will be highlight in **red** in Table 5.

> **Note that** precursor ions were binned by 10 Da for Pearson correlation calculation.

Table 5: Precursor Distribution across mass similarity

|  | QC\_p (1).mzML | QC\_p (10).mzML | QC\_p (2).mzML | QC\_p (3).mzML | QC\_p (4).mzML | QC\_p (5).mzML | QC\_p (6).mzML | QC\_p (7).mzML | QC\_p (8).mzML | QC\_p (9).mzML |
| --- | --- | --- | --- | --- | --- | --- | --- | --- | --- | --- |
| QC\_p (1).mzML | 1 | 0.989 | 0.997 | 0.999 | 0.997 | 0.994 | 0.991 | 0.984 | 0.991 | 0.989 |
| QC\_p (10).mzML | 0.989 | 1 | 0.993 | 0.993 | 0.991 | 0.994 | 0.996 | 0.994 | 0.997 | 0.998 |
| QC\_p (2).mzML | 0.997 | 0.993 | 1 | 0.997 | 0.997 | 0.995 | 0.993 | 0.989 | 0.993 | 0.993 |
| QC\_p (3).mzML | 0.999 | 0.993 | 0.997 | 1 | 0.998 | 0.998 | 0.996 | 0.988 | 0.994 | 0.993 |
| QC\_p (4).mzML | 0.997 | 0.991 | 0.997 | 0.998 | 1 | 0.997 | 0.994 | 0.985 | 0.991 | 0.99 |
| QC\_p (5).mzML | 0.994 | 0.994 | 0.995 | 0.998 | 0.997 | 1 | 0.999 | 0.987 | 0.993 | 0.993 |
| QC\_p (6).mzML | 0.991 | 0.996 | 0.993 | 0.996 | 0.994 | 0.999 | 1 | 0.99 | 0.995 | 0.996 |
| QC\_p (7).mzML | 0.984 | 0.994 | 0.989 | 0.988 | 0.985 | 0.987 | 0.99 | 1 | 0.997 | 0.997 |
| QC\_p (8).mzML | 0.991 | 0.997 | 0.993 | 0.994 | 0.991 | 0.993 | 0.995 | 0.997 | 1 | 1 |
| QC\_p (9).mzML | 0.989 | 0.998 | 0.993 | 0.993 | 0.99 | 0.993 | 0.996 | 0.997 | 1 | 1 |

#### 4.3 Precursor Distribution across RT

`Precursor Distribution across RT` plot is used to visualize the number of fragmented precursors at each RT (or scan). It can be useful to spot any signal dropouts during data acquisition.

- In both **data-dependent acquisition (DDA)** and **data-independent acquisition (DIA)** mode, you are expected to see a continuous precursor ions distribution across the entire RT range.
- High similarity in `Precursor Distribution across RT` among QC samples indicates good reproducibility in MS2.

```
## $`1`
```

```
## 
## $`2`
```

`Cosine Similarity` is used to quantify the similarity of precursor distribution across RT plots among QC samples. Cosine similarity over 0.85 indicate high similarity. If the value is below 0.85, it will be highlight in **red** in Table 6.

> **Note that** RT were binned by 0.05 min for Pearson correlation calculation.

Table 6: Precursor Distribution across RT similarity

|  | QC\_p..1..mzML | QC\_p..10..mzML | QC\_p..2..mzML | QC\_p..3..mzML | QC\_p..4..mzML | QC\_p..5..mzML | QC\_p..6..mzML | QC\_p..7..mzML | QC\_p..8..mzML | QC\_p..9..mzML |
| --- | --- | --- | --- | --- | --- | --- | --- | --- | --- | --- |
| QC\_p..1..mzML | 1 | 0.998 | 0.998 | 0.998 | 0.999 | 0.999 | 0.999 | 0.993 | 0.996 | 0.996 |
| QC\_p..10..mzML | 0.998 | 1 | 0.997 | 0.998 | 0.998 | 0.998 | 0.998 | 0.996 | 0.997 | 0.997 |
| QC\_p..2..mzML | 0.998 | 0.997 | 1 | 0.998 | 0.999 | 0.999 | 0.998 | 0.992 | 0.995 | 0.995 |
| QC\_p..3..mzML | 0.998 | 0.998 | 0.998 | 1 | 0.999 | 0.999 | 0.998 | 0.994 | 0.996 | 0.996 |
| QC\_p..4..mzML | 0.999 | 0.998 | 0.999 | 0.999 | 1 | 0.999 | 0.998 | 0.993 | 0.996 | 0.996 |
| QC\_p..5..mzML | 0.999 | 0.998 | 0.999 | 0.999 | 0.999 | 1 | 0.998 | 0.994 | 0.996 | 0.996 |
| QC\_p..6..mzML | 0.999 | 0.998 | 0.998 | 0.998 | 0.998 | 0.998 | 1 | 0.994 | 0.997 | 0.997 |
| QC\_p..7..mzML | 0.993 | 0.996 | 0.992 | 0.994 | 0.993 | 0.994 | 0.994 | 1 | 0.997 | 0.997 |
| QC\_p..8..mzML | 0.996 | 0.997 | 0.995 | 0.996 | 0.996 | 0.996 | 0.997 | 0.997 | 1 | 0.998 |
| QC\_p..9..mzML | 0.996 | 0.997 | 0.995 | 0.996 | 0.996 | 0.996 | 0.997 | 0.997 | 0.998 | 1 |
